# A Variable Clock Underlies Internally Generated Hippocampal Sequences

**DOI:** 10.1101/2020.04.10.035980

**Authors:** Xinyi Deng, Shizhe Chen, Marielena Sosa, Mattias P. Karlsson, Xue-Xin Wei, Loren M. Frank

**Author notes:** **For correspondence:** (XD); (LMF).

## Abstract

Humans have the ability to store and retrieve memories with various degrees of specificity, and recent advances in reinforcement learning have identified benefits to learning when past experience is represented at different levels of temporal abstraction. How this flexibility might be implemented in the brain remains unclear. We analyzed the temporal organization of rat hippocampal population spiking to identify potential substrates for temporally flexible representations. We examined activity both during locomotion and during memory-associated population events known as sharp wave-ripples (SWRs). We found that spiking during SWRs is rhythmically organized with higher event-to-event variability than spiking during locomotion-associated population events. Decoding analyses using clusterless methods further indicate that similar spatial experience can be replayed in multiple SWRs, each time with a different rhythmic structure whose periodicity is sampled from a lognormal distribution. This variability increases with experience despite the decline in SWR rates that occurs as environments become more familiar. We hypothesize that the variability in temporal organization of hippocampal spiking provides a mechanism for storing experiences with various degrees of specificity.

---

The human brain has the remarkable ability to call up memories with different levels of temporal specificity, including experiences that range in extent from seconds to perhaps years. The potential utility of this ability has been demonstrated by a recently proposed theoretical reinforcement learning framework (***Levy et al***., 2019). This work showed that hierarchical agents that operate at different levels of temporal abstraction by using “multilevel hindsight experience replay” can learn tasks more quickly, both because they can divide the work of learning behaviors among multiple policies and because they can also explore internal representations of the environment at varying levels of temporal resolution. This result highlights the utility of variable temporal abstraction in the theoretical framework of deep reinforcement learning (***Schmidhuber***, 1992; ***Sutton et al***., 1999; ***Bakker and Schmidhuber***, 2004; ***Kulkarni et al***., 2016; ***Konidaris***, 2019). How the brain might implement this flexibility remains unknown.

A potential mechanism for this flexibility would involve storing experiences with different levels of temporal specificity to facilitate subseqeunt retrieval at different levels of specificity. To determine whether the brain might engage such a mechanism, we can examine activity patterns in the rodent hippocampus, a structure critical for storing and retrieving memories for spatial experiences (***Eichenbaum and Cohen***, 2004). As an animal moves through its environment, individual hippocampal neurons (place cells) are active when the animal occupies specific regions of space, known as the cells’ place fields (***O’Keefe and Dostrovsky***, 1971). In the context of this “on-line” state (***Buzsaki***, 1989, 2002, 2019; ***Kay and Frank***, 2019), a traversal through a given environment results in the sequential activation of a series of these place cells. This sequential activation is modulated in association with the ~8 Hz theta oscillation, with the result that firing is most prevalent at the trough and least prevalent at the peak of each theta cycle (***Buzsaki***, 2002). Importantly, the frequency of theta varies across a range of only ~1 Hz, with higher frequencies associated with higher running speeds (***Hinman et al***., 2011). Thus, theta can be understood as stable internal clocking mechanism that organizes spiking activity in the hippocampal network (***Buzsaki***, 2002).

Hippocampal spiking is also organized within each theta cycle. Gamma frequency (~40 Hz) oscillations often occur together with theta oscillations in the hippocampal LFP and these gamma oscillations modulate hippocampal spiking as well (***Bragin et al***., 1995; ***Lisman and Buzsaki***, 2008; ***Colgin et al***., 2009; ***Kemere et al***., 2013; ***Lisman and Jensen***, 2013; ***Lopes-Dos-Santos et al***., 2018). This modulation has been proposed to help maintain a further level of organization of spiking within each theta cycle that is known as a “theta sequence” (***Lisman***, 2005): within each cycle, sets of cells with overlapping fields will typically fire in a compressed sequence that often recapitulates the longer timescale sequential activity associated with the serial order of the place fields (***Skaggs et al***., 1996; ***Foster and Wilson***, 2006; ***Colgin et al***., 2009; ***Gupta et al***., 2012).

Time-compressed versions of these sequences are also seen during hippocampal Sharp-Wave Ripple (SWR) events (***Foster and Wilson***, 2006; ***Lee and Wilson***, 2002; ***Karlsson and Frank***, 2009; ***Davidson et al***., 2009). This replay serves as a mechanism for the reinstatement of previously stored representations (e.g. retrieval) that is thought to promote the longer term storage and updating of these representations in distributed cortical networks (consolidation) (***Buzsaki***, 2015; ***Joo and Frank***, 2018; ***Gillespie et al***., 2021).

Could SWRs support the storage of memories in a way that would allow for temporally flexible retrieval? If so, then we might expect that the temporal structure of these events might vary from event to event. SWRs are named in part because of the associated high frequency (150-250 Hz) ripple oscillation, but this oscillation is generated locally in hippocampal area CA1 and does not coordinate the replay of experience across the hippocampal network (***Carr et al***., 2012; ***Sirota et al***., 2003). This coordination may instead be mediated by a slower process. Consistent with that possibility, during SWRs there are transient increases in slow gamma (20-50 Hz) power and synchrony across dorsal CA3 and CA1 networks of both hemispheres (***Carr et al***., 2012; ***Gillespie et al***., 2016; ***Oliva et al***., 2018). Slow gamma phase also provides a reliable “clock” for replay events (***Carr et al***., 2012), and it has been suggested that these gamma oscillations govern temporal segmentation of spatial content in replay sequences: during phases of high neural activity within the gamma cycle, spatial representation is often focused on a single location, whereas during phases of low neural activity, the spatial representation is more likely to move to adjacent locations (***Pfeiffer and Foster***, 2015).

Slow gamma varies over a wide range, but precisely how this variation manifests in hippocampal network during SWRs has not been examined. Moreover, whether this variability is related to the representational content of SWRs is also unknown. To address these issues, we examined the temporal organization of population spiking events on an event-to-event basis during locomotion and awake immobility as rats learned to perform a memory-guided task. We found that population spiking underlying SWR events was rhythmic and exhibited much higher event-to-event variability in their periodicity than locomotion-associated spiking sequences. Events with varying periodicity could represent similar spatial experiences, and, surprisingly, variability increased rather than decreased as the environment became more familiar. We hypothesize that this experience-dependent variability in SWR rhythmic organization supports memory storage in support of temporally flexible memory retrieval.

## RESULTS

### Population Spiking Underlying Event-to-Event Variability

To examine the temporal organization of hippocampal population spiking, we analyzed previously recorded electrophysiological data from four male Long-Evans rats implanted with a microdrive array containing multiple independently movable tetrodes targeting CA1 and CA3 (***Karlsson and Frank***, 2008, 2009; ***Kay et al***., 2016). Data were collected as animals learned to perform a memory-guided alternation task in a W-shaped environment (**Figure 1A**). The animal was rewarded each time it visited the end of an arm in the correct sequence, starting in the center and then alternating visits to each outer arm and returning to the center (see **Materials and Methods**).

**Figure 1.**
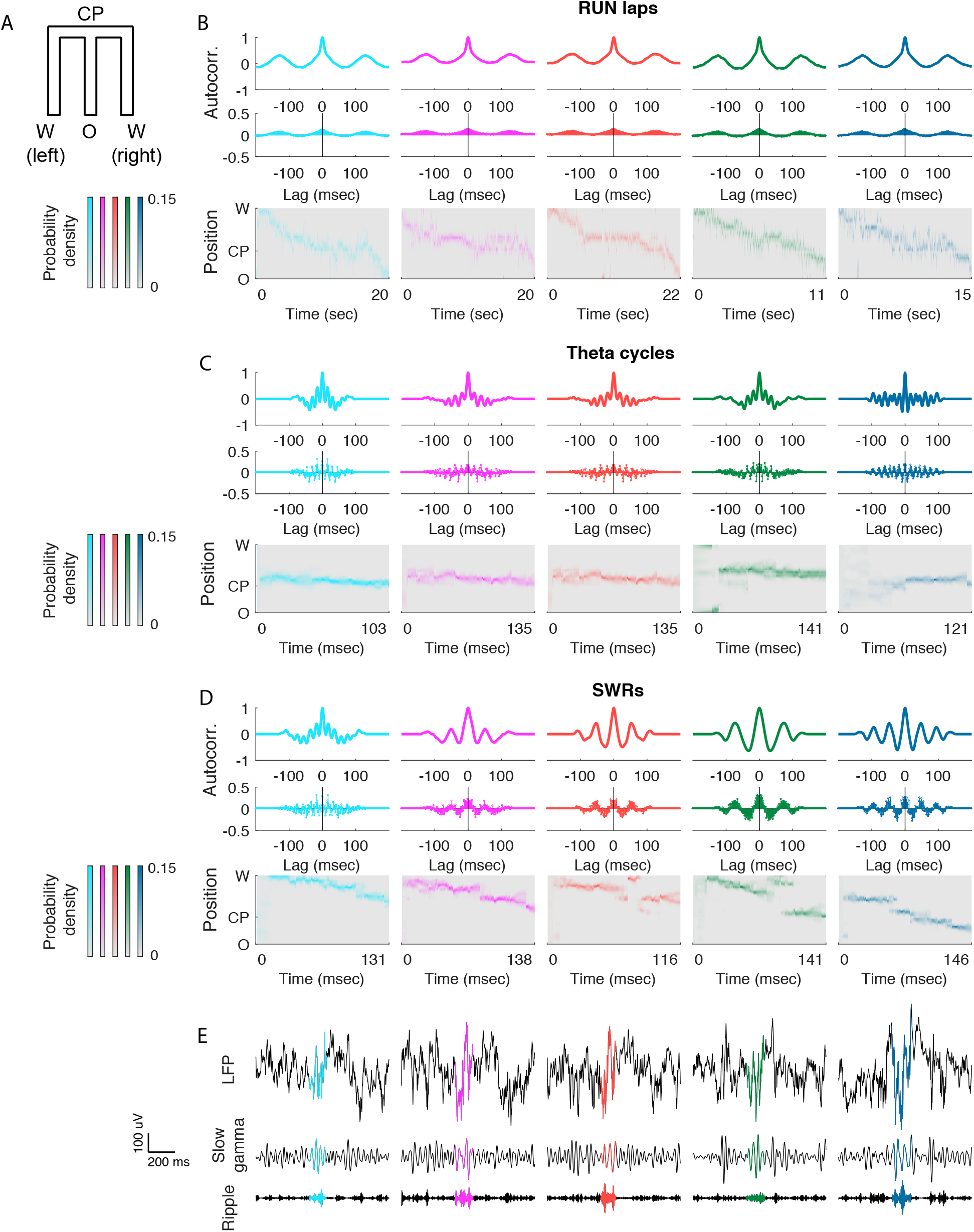
SWR-associated temporal organization of population spiking is rhythmic and exhibits higher event-to-event variability than locomotion-associated population spiking. (**A**) 2D representation of an W-maze. (**B**) Rhythmicity of spiking across traversals of an inbound (W→O) trajectory in the W-maze (RUN laps) from Animal 2. Each column represents one RUN lap. *Top*, peak-normalized Gaussian-smoothed autocorrelation function of population spiking. *Middle*, autocorrelation function of population spiking. *Bottom*, decoded trajectory where the heat plot shows the estimated posterior density at each time step. Theta (~125 ms) modulation of autocorrelation is clearly visible across different trajectories. (**C**) Rhythmicity of spiking across theta cycles, rows as in (**B**), taken from inbound runs on Animal 2. Each column represents one theta cycle. Population spiking within each theta cycle was also rhythmically organized with a periodicity within the gamma range (mean period ~18 ms). (**D**) Rhythmicity across SWR events, rows as in (**B**), with SWRs from an example epoch of Animal 2 with inbound spatial content. Each column represents one SWR with inbound representational content. (**E**) Raw and filtered local field potential (LFP) signals from one CA1 tetrode for corresponding SWR event in Figure 1(**D**). *Top*, raw LFP trace; *Middle*, slow gamma band 20-50 Hz; *Bottom*, ripple band 150-250 Hz. The colored highlights of each LFP trace denote the entire duration of an individual SWR event.

Our first goal was to identify regularities in timing of spiking within each individual population spiking event during locomotion and awake immobility. The standard approach, using “clustered” spikes that have been assigned to individual single units, discards the much larger number of spikes that cannot be confidently assigned to a single unit, and the resulting sparse data may not be sufficient for inferring temporal structure. As such, we used the unclustered multiunit spiking data collected across all CA1 and CA3 tetrodes for our analyses (***Kloosterman et al***., 2014; ***Deng et al***., 2015).

The rhythmicity of the hippocampal spiking activity during a specific period of time was determined by calculating the autocorrelation function of the multiunit spiking activity. Specifically, the “clock speed” for each event was defined as the lag of the first positive side peak of the autocorrelation, reflecting the average time between spiking events. Note that clock speed and timing of the first positive peak lag of the SWR autocorrelation function are inversely correlated, with lower value of peak lag denoting fasting clock speed. Spike analyses with autocorrelation also avoids the usual caveat of superposition of oscillations with different phases in LFP analyses. In parallel, we decoded the spatial representation expressed during those times using a clusterless decoding method (see **Materials and Methods**; ***Deng et al***., 2015) that does not require multiunit spiking waveforms to be sorted into single units and instead incorporates waveform information from all spikes.

We first computed the autocorrelation function of hippocampal population spiking during individual locomotion laps (**Figure 1B** shows five example inbound laps) and as expected, the autocorrelation function had peak at the stereotypical theta lag of ~125 ms. We then computed the autocorrelation function of hippocampal population spiking during individual theta cycles (defined by a consensus measure between LFP and multiunit spiking; see **Materials and Methods**). Once again as expected, population activity within each theta cycle was also rhythmically organized with a periodicity within the gamma range (**Figure 1C**; ***Lisman***, 2005; ***Lisman and Buzsaki***, 2008; ***Colgin et al***., 2009; ***Lisman and Jensen***, 2013; ***Lopes-Dos-Santos et al***., 2018).

Finally, we computed the autocorrelation function of hippocampal population spiking during individual SWR events. We observed that population spiking activity during individual SWRs is strikingly rhythmically modulated (**Figure 1D**, LFP traces in **Figure 1E**), consistent with previous demonstrations that SWR events include slow gamma oscillations (***Carr et al***., 2012; ***Pfeiffer and Foster***, 2015). Interestingly, the SWR-associated spiking organization appeared to exhibit very high event-to-event variability, even across replay events that express similar spatial representations.

### Periodicity of SWR-associated Rhythmic Organization Follows an Approximate Lognormal Distribution

To evaluate whether the rhythmic spiking organization across both locomotion-associated and SWR-associated hippocampal population events varies continuously or clusters into discrete bands of frequency, we collated the autocorrelation functions of spiking activity of all hippocampal spiking events within each recording epoch and ordered the events by the timings of the first positive peak lag of the autocorrelation function (**Figure 2A**).

**Figure 2.**
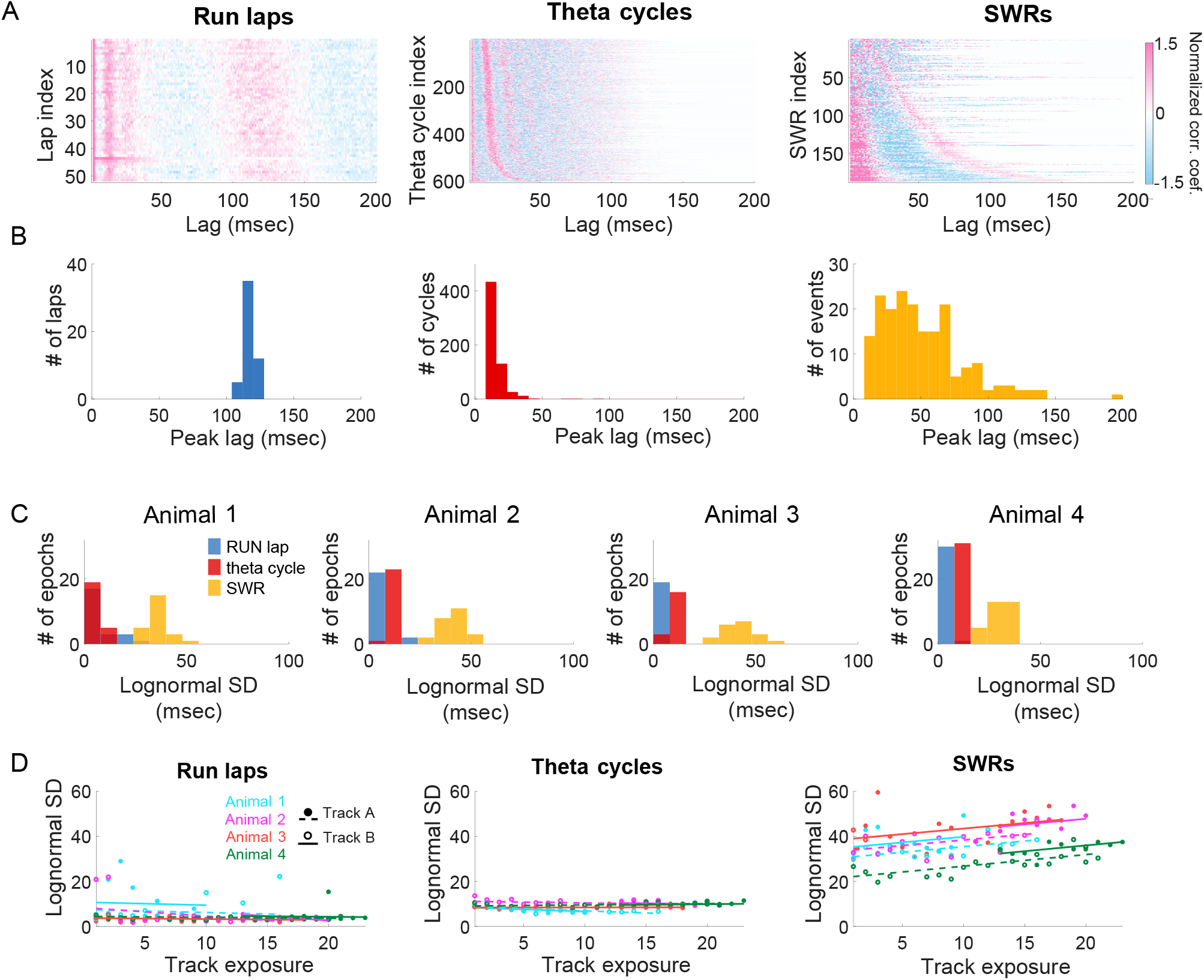
Lognormal distribution of periodicity for SWR-associated spiking. (**A, B**) Periodicity of the rhythmic spiking organization underlying RUN laps clusters tightly around 125 ms (*left*), periodicity for theta cycles clusters tightly around 15 ms (*middle*), while periodicity for SWRs spans a continuous range between 20 ms and 160 ms (*right*). (**A**) Normalized autocorrelations of multiunit spiking activity during individual RUN laps (*left*), theta cycles (*middle*), and SWR events (*right*) of an example epoch of Animal 3, collated in ascending order by the timings of the first positive side peak lag. Every horizontal line is the autocorrelation function of an individual lap, theta cycle, or SWR, respectively; only non-negative lags are displayed. Positive and negative correlation coefficients are plotted in shades of red and blue, respectively. (**B**) Histogram of the timings of the first positive side peak lag of the multiunit spiking autocorrelation lags across RUN laps (*left*), theta cycles (*middle*), or SWR events (*right*), respectively, for an example epoch. (**C**) Periodicity of SWR-associated spiking organization follows an approximate lognormal distribution. Each panel plots the histogram of the standard deviations of lognormal distribution fit to peak lag distributions across all epochs for an animal. (**D**) Standard deviation of lognormal distribution fit to SWR-associated spiking organization positively correlates with track exposure. Relationship between the number of exposures to a track environment and the standard deviation (SD) of lognormal distribution fit to peak lag distributions during RUN laps (*left*), theta cycles (*middle*), and SWR events (*right*). Each color denotes an animal. Each filled circle represents one epoch recorded in track A, and each empty circles represents one epoch recorded in track B.

We found that the periodicity of the rhythmic spiking organization underlying locomotion (RUN) laps clusters tightly at ~125 ms, consistent with ~8Hz theta (Figure 2B, *left panel*), while periodicity for theta cycles clusters tightly at ~18 ms, consistent with the high end of the typical ~20-55 Hz slow gamma range (**Figure 2B**, *middle panel*), respectively. We also observed that when grouped by movement speed, theta cycles with lower speeds were more likely to be associated with larger mode values of the empirical distributions (**Figure S1**; mixed effects model, p < 0.0001), consistent with an increased prevalence of slow gamma at lower movement speeds (***Kemere et al***., 2013).

In contrast, the periodicity of the rhythmic SWR-associated population spiking spanned a continuous range between ~6Hz and ~50Hz (**Figure 2B**, *right panel*). This continuous distribution was seen across the multiple run sessions within each day, across days, and across animals (**Figure S2**). The distribution was also visible in both areas CA1 and CA3, although the organization was clearer in CA1, perhaps due to the larger proportions of active neurons in CA1 as compared to CA3 (**Figure S3**, *top panels*; ***Karlsson and Frank***, 2008). The distribution was also visible across both putative excitatory and inhibitory cell types (**Figure S3**, *bottom panels*).

The periodicities underlying SWRs spanned a much wider range than those underlying RUN laps or theta cycles. To quantify this difference in distributional spread, we fit parametric distributions to each empirical distribution (**Figure S4**). We found that the periodicities of SWR events were approximately lognormal, with the distribution median at the low end of the slow gamma range: exp(mu) =48 ms. Such a lognormal distribution creates a periodicity spectrum with a wide dynamic range, spanning from the majority of SWR events with a period between 25 ms and 75 ms to a small fraction of events with either a slower or a faster rhythmic organization. Importantly, the SWR-associated lognormal distributions had significantly larger standard deviation than locomotion-associated lognormal distributions (**Figure 2C**; across all four animals, p < 0.001, bootstrap tests).

If this broad range of clock speeds serves an important function, it should be preserved throughout experiences in a given environment. Previous work has demonstrated that novel experiences drive high SWRs rates (***O’Neill et al***., 2006; ***Cheng and Frank***, 2008), which then fall by about half as the environment becomes more familiar. To determine whether the range of clock speeds is preserved as the environment becomes more familiar and SWR rates decrease, we constructed a linear mixed-effects model (random intercept and random slope for exposure with animal-specific random effects). This model captures the modulation of the standard deviation of the lognormal distribution by the number of exposures to each track environment (**Figure 2D**; for empirial distributions, see **Figure S5**). Strikingly, we found that the standard deviation that characterizes the distributional spread increased with familiarity. The standard deviation was consistently and significantly positively correlated with the number of track exposure, for SWRs (across four animals, p < 10^−6^, t-test for fixed effects), but not for locomotion-associated spiking events (across four animals, p = 0.25 for RUN laps, p = 0.30 for theta cycles, t-test for fixed effects; median of empirical SWR event durations and proportion of SWR duration longer than 100 msec showed no consistent trend across track environment nor across animals, see **Figure S6**). For visualization purposes, we plotted the regression lines for each track environment for each animal and observed that in the case of SWRs, they were approximately parallel to each other (**Figure 2D**, *right panel*). This indicates a consistent change across environments and across animals, where the lognormal distribution of SWRs, but not locomotion-associated events, becomes broader as the environment becomes more familiar.

### Periodicity of SWR-associated Rhythmic Organization Is Correlated with LFP Ripple Power

How does this continuously distributed “clock” relate to the LFP? To address this question, we grouped all detected SWR events in a recording epoch into five categories by the timings of the first positive peak lag of the SWR autocorrelation functions, from events with a fast rhythmic organization with periodicity less than 25 ms to events with a stereotypical slow gamma organization with periodicity between 25 ms and 50 ms, to events with a slow rhythmic organization with periodicity more than 100 ms. We then computed the average SWR-triggered spectrum of the LFP, normalized by the baseline spectrogram across an entire recording epoch (see **Materials and Methods**) for each of the five groups of SWR events (**Figure 3A**). We observed that the strongest concentration of power in the slow gamma band emerges in the groups whose autocorrelation lags have peaks between 25 ms and 75 ms. This concentration of power gradually shifts to lower frequencies as the peak lag of the SWR autocorrelation increases. We also observed increased power at ripple band as the peak lag of the autocorrelation increases.

**Figure 3.**
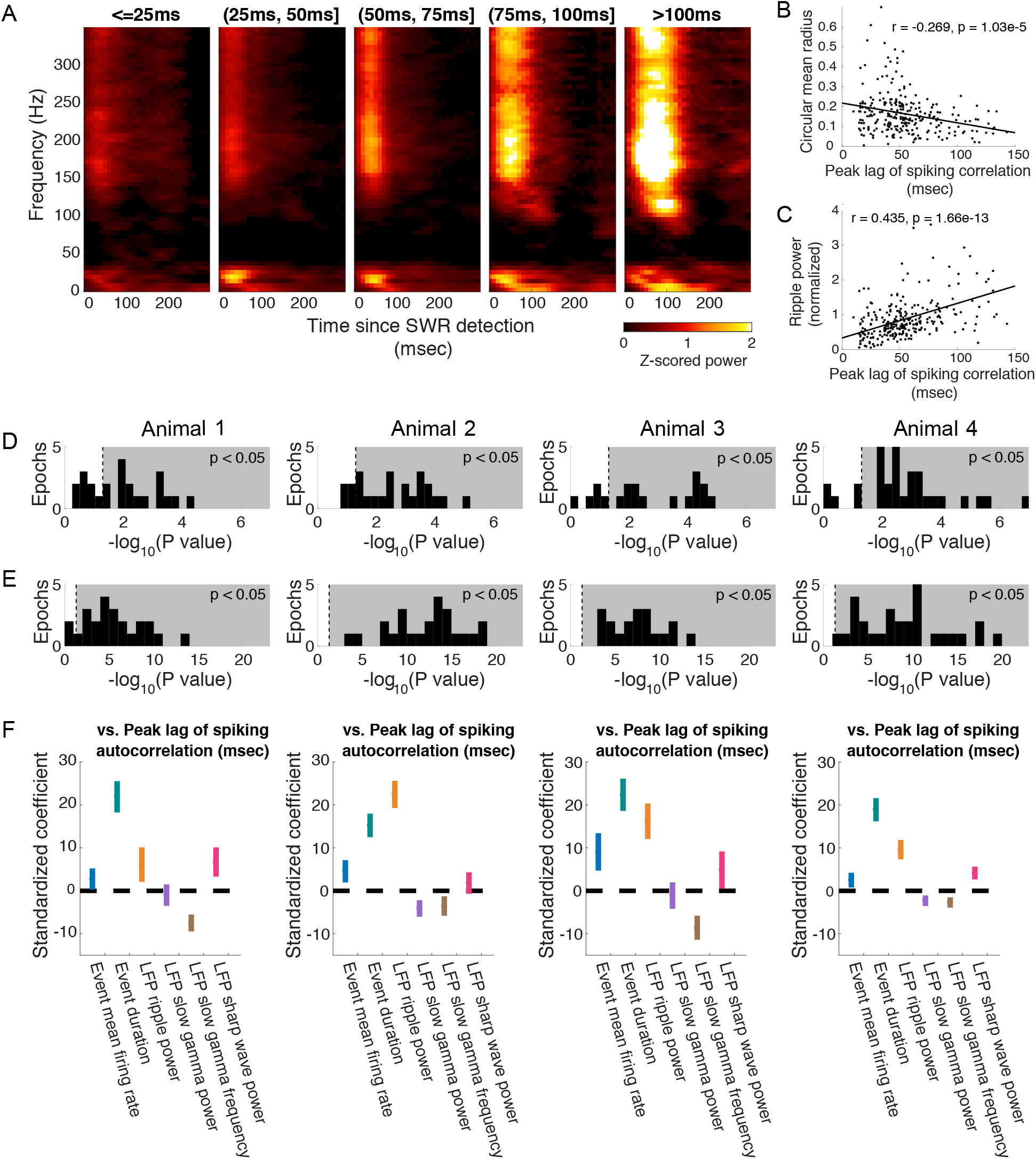
SWR-associated spiking organization correlates with strength of gamma phase locking and ripple power in local field. (**A**) Average SWR-triggered LFP spectrograms grouped by first positive autocorrelation side peak (peak lag of autocorrelation) for an example epoch for Animal 2. (**B**) Negative correlation between gamma phase locking (circular mean radius) of multiunit activity and peak lag for an example epoch for Animal 2. Insert, r, Pearson correlation coefficient; p, p-value fortesting hypothesis of no correlation. (**C**) Positive correlation between normalized ripple power and peak lag for an example epoch for Animal 2. Insert, r, Pearson correlation coefficient; p, p-value for testing hypothesis of no correlation. (**D**) Histogram, for each animal, of −log(p-value) for the Pearson correlation coefficient between strength of the gamma phase locking of multiunit activity and peak lag of the SWR autocorrelation. Grey background indicates the area with p < 0.05. (**E**) Histogram, for each animal, of −log(p-value) for the Pearson correlation coefficient between normalized LFP ripple power and peak lag of the SWR autocorrelation. Grey background indicates the area with p < 0.05. (**F**) Regression coefficients and confidence intervals for six SWR-associated covariates as regressors for peak lag for each animal. Vertical line segments indicate 95% confidence intervals of the standardized coefficients for fixed effects of event mean firing rate (plotted in blue), event duration (green), LFP ripple power (yellow), LFP slow gamma power (purple), LFP slow gamma frequency (brown), and LFP sharp wave power (red) in a multiple linear mixed-effects regression model that allows random effects for epochs. Consistent across animals, duration of SWR events and LFP ripple power were significantly positively correlated with the peak lag of the spiking autocorrelation, while LFP slow gamma frequency was significantly negatively correlated with the peak lag of the spiking autocorrelation.

We then quantified these relationships. We first measured the strength of the slow gamma phase locking of the multiunit spiking activity and confirmed that across all lags, there was significant phase locking to slow gamma (t-test, p < 10^−12^; across all four animals, t-tests for 100% of epochs, p < 0.001). We also observed that slow gamma phase locking decreased as the clock speed increased (Pearson’s r = −0.269, p < 10^−4^, **Figure 3B**; for analyses across all four animals, see **Figure 3D**). Importantly, this phase locking was also present for CA3 spikes alone (across all four animals, p< 10^−12^, Rayleigh’s test for detecting a unimodal deviation from circular uniformity), as expected given previous results (***Carr et al***., 2012). In contrast, there was a significant inverse relationship between ripple power and clock speed, with lower normalized ripple power at higher clock speed (Pearson’s r = 0.435, p < 10^−12^, **Figure 3C**; for analyses across all four animals, see **Figure 3E**).

These initial analyses identified covariates of the clock speed, and motivated a more comprehensive model. We therefore constructed a multiple linear mixed-effects regression model with random effects of epochs that captured the modulation of the SWR-associated spiking organization with a set of parameters defining the influence of mean population spiking rate, SWR event duration, LFP ripple power, LFP slow gamma power, and LFP slow gamma peak frequency (Figure 3F). We estimated the values of these parameters that maximize the likelihood of the observed SWR population spiking. The regression analysis showed that LFP slow gamma frequency was significantly negatively correlated with the peak lag of the spiking autocorrelation (across all epochs for four animals, p < 0.01, t-tests for fixed effects), which is consistent with the qualitative observations from the previous spectrogram analysis (Figure 3A). Population firing rate was positively correlated with the peak lag of the spiking autocorrelation at the significance level of 0.01 in one of the four animals (p = 2.675 × 10^5^, t-test for fixed effects), at the significance level of 0.05 in another animal (p = 0.0486, t-test for fixed effects), but not in the other two animals (p = 0.4088 and p = 0.1277, t-tests for fixed effects). We also found that the duration of individual SWR events was significantly positively correlated with the peak lag of the spiking autocorrelation (across all epochs for four animals, p< 10^−5^, t-tests for fixed effects). Thus, as one might expect, the slower the hippocampal rhythmic organization underlying an individual SWR event is, the longer the duration of the event is. The regression analysis further showed that, after accounting for effects of other SWR temporal and frequency components, LFP ripple power remained significantly positively correlated with the peak lag of the spiking autocorrelation (across epochs for four animals, p < 0.01, t-tests for fixed effects).

### Periodicity of SWR-associated Rhythmic Organization Is Correlated with Classifiability, but Not Types, Of Representational Content

Finally, we asked whether the rhythmic spiking organization of an SWR event relate to its representational content. We applied a previously developed and validated discrete decision state point process filter (see **Materials and Methods**; ***Deng et al***., 2016) to classify the representational content of SWR events from multiunit spiking activity into four categories: an “outbound, forward” path, an “outbound, reverse” path, an “inbound, forward” path, or an “inbound, reverse”, based on the direction and temporal order of the spatial trajectory and computed our confidence about the classification; events whose decision state probabilities did not pass the confidence threshold and cannot be classified into any of the four aforementioned categories are denoted as “unclassified” (see **Figure 4A** for example SWRs).

**Figure 4.**
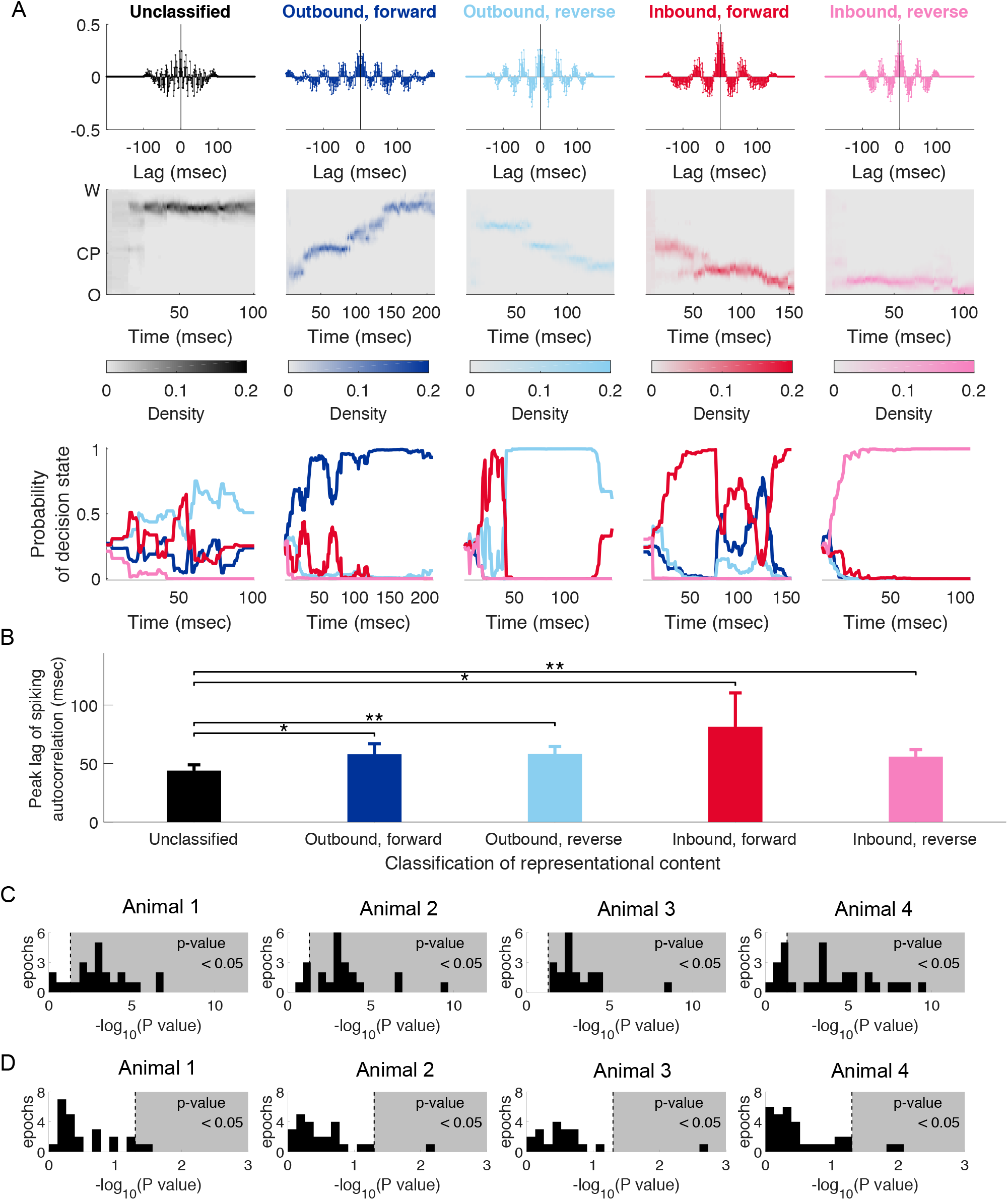
SWR-associated spiking organization correlates with classifiability of representational content, but not the type of the content. (**A**) Examples of SWR events with particular classifications of representational content. Each column represents one SWR. *Top*, autocorrelation function of population spiking. *Middle*, Decoded trajectory where the heat plot shows the estimated posterior density at each time step. *Bottom*, Probability of the decision state as a function of time. Probability of the SWR event representing an “outbound, forward” path, an “outbound, reverse” path, an “inbound, forward” path, or an “inbound, reverse” path is plotted in darker blue, lighter blue, darker red, and lighter red, respectively. (**B**) Peak lags of the SWR autocorrelation function grouped by classification of representational content for an example epoch of Animal 2. The group of SWR events whose decision state probabilities did not pass the threshold and cannot be classified is plotted in gray. The groups of SWR events whose representational content can be classified with confidence as an “outbound, forward” path, an “outbound, reverse” path, an “inbound, forward” path, or an “inbound, reverse” path are plotted in darker blue, lighter blue, darker red, and lighter red, respectively. All bar graphs show mean + 1.96 × SEM. Lines and asterisks indicating significant differences between pairs of groups under Wilcoxon rank-sum one-sided tests with Bonferroni correction. * p-value < 0.05; ** p-value < 0.01. (**C**) Histogram, for each animal, of p-value of Wilcoxon rank-sum tests for an ordinal difference between the peak lags of the spiking autocorrelations of the unclassified events and that of the classified events across epochs. (**D**) Histogram, for each animal, of p-values of Kruskal-Wallis tests for a difference between the peak lags of the spiking autocorrelations across groups with different types of representational content among classified events across epochs.

When we examined all SWRs, we found there was a statistically significant difference between the peak lag of the spiking autocorrelation of the unclassified group and that of each of the classified groups (see **Figure 4B** for an example epoch: Wilcoxon rank-sum one-sided tests with Bonferroni correction, p(“unclassified” vs. “outbound, forward”) = 0.035, p(“unclassified” vs. “outbound, reverse”) = 0.0078, p(“unclassified” vs.”inbound, reverse”) = 0.021, and p(“unclassified” vs. “inbound, reverse”) = 0.0027; for analyses across all four animals, see **Figure 4C**). This result suggests that SWRs with unclassified content tend to have a faster rhythmic organization while SWRs with classifiable content tend to have a slower one. In other words, the periodicity of the rhythmic structure underlying an SWR is related to the classifiability of its representational content. We then examined only SWRs with classifiable content and found no statistically significant differences (Kruskal-Wallis tests; for analyses across all four animals, see **Figure 4D**). Thus, among SWRs with classifiable content, there is no correlation between periodicity of rhythmic spiking organization and the type of content.

## DISCUSSION

We explored the structure of spiking activity across periods of locomotion and during hippocampal sharp-wave ripple (SWR) events. We found that the periodicity of the SWR-associated hippocampal rhythmic spiking organization has much higher event-to-event variability than locomotion-associated spiking. This SWR-associated clocking variability was observed across recording epochs, across experimental days, and across animals. The wide range of “clocks” was clearly visible in novel environments where SWRs are very prevalent (***O’Neill et al***., 2006; ***Cheng and Frank***, 2008) and actually increased in more familiar environments. No such changes were present for locomotion-associated spiking. The large variability in the temporal organization of SWRs, combined with the observation that the brain can maintain less variable timing during locomotion, demonstrates that lower variability is within the capacities of the system, and suggests that the higher variability seen during SWRs serves a function.

### SWR periodicity and slow gamma

We also found that the periodicity of this “clock” is related to the frequency of the slow gamma rhythm in each SWR. This is consistent with prior work (***Carr et al***., 2012; ***Pfeiffer and Foster***, 2015) and also with the results presented in a recent report from Oliva and colleagues (***Oliva et al***., 2018), although those authors came to a different conclusion about the data. They claimed that the slow gamma power increase underlying SWRs is a “spurious” oscillation because “slow gamma power is specifically associated with longer SPW-Rs produced by the overlap of multiple ripple events” (***Oliva et al***., 2018). This conclusion was partially based on a replication of the observation of coupling between slow-gamma phase and ripple amplitude (***Carr et al***., 2012), indicating bursts of ripple power at slow gamma frequency. The authors did not present a definition of an actual oscillation that would allow for a clear separation from spurious oscillations however, making it difficult to evaluate their conclusion. Their conclusion is also puzzling given the well-established presence of slow gamma as a reflection of CA3 input to CA1 (***Colgin et al***., 2009; ***Kemere et al***., 2013), the locking of CA3 and CA1 spikes to slow gamma during SWRS (***Carr et al***., 2012), and the known role of CA3 input in driving SWRs in CA1 (***Buzsaki***, 2015).

We suggest instead that the question should be whether an oscillation extracted from the wideband LFP reflects a real physiological process (in this case CA3 input to CA1) and is useful in understanding information processing in the brain. Using that criterion, our findings corroborate previous work (***Carr et al***., 2012; ***Pfeiffer and Foster***, 2015) and indicate that measuring power and phase in the slow gamma provides important information about SWR characteristics, including the presence of sequential content (***Carr et al***., 2012). We found that, despite of the wide range of clock speeds resulting in different numbers of clock cycles for each individual SWR event, multiunit spiking in CA3 was significantly phase-clocked to the slow gamma clock in CA1. We further showed that, even after accounting for event duration, mean firing rate, and other local field frequency components, the periodicity of this slower, rhythmic spiking process that underlies SWR events remains significantly correlated with LFP ripple power, which relates to pyramidal cell synchrony (***Csicsvari et al***., 2000). These findings strengthen the association between the temporal structure of SWRs and the robustness of spiking activity within the events. We therefore suggest that population synchrony in the hippocampal circuits mediated by a slow gamma “clock” is more robust, thus leads to longer SWR events. At the same time, we identified a small number of events with a “clock” as slow as ~6Hz. This suggests that the slow gamma range may better be defined as a lognormal distribution that extends to lower and higher frequencies.

### A Changing Hippocampal “Clock” Is a Potential Mechanism for Flexible Memory Retrieval

Is there a possible functional role of the SWR-associated rhythmic organization and of its variability? There are two possibilities. One is that the brain wishes to impose a uniform temporal structure when storing or updating representations but due to intrinsic variability (***Atallah and Scanziani***, 2009; ***Hemberger et al***., 2019), it fails to do so and produces variably timed replay events. The other possibility is that replaying the same spatial content with varying degree of rhythmic organization is beneficial for flexible storage. In this scenario, the goal would be to store memories with a more flexible temporal structure that would provide for different degrees of “chunking” in the hippocampus and, presumably, in down-stream cortical networks that are engaged during SWRs (***Buzsaki et al***., 2013).

Our data are more consistent with the second of these two possibilities. We found that the hippocampus can express sequences with very low temporal variability, both across and within theta cycles seen during locomotion. This indicates that the hippocampus can generate spiking patterns that are tightly locked to a given frequency. By contrast, the temporal variability of SWR-related spiking is high during new experiences, and increases as the environment becomes more familiar. In other words, as an animal learns the task in an environment, the hippocampal circuits replay its spatial experience with an underlying “clock” sampled from an increasingly broader lognormal distribution. We note that in our dataset, this is not associated with a consistent experience-dependent trend for its median or tail probability (proportion of event longer than 100 msec) (***Fernandez-Ruiz et al***., 2019).

This increasing variability indicates that the variability in the “clock” is not solely a result of instability associated with recent learning, and suggests that the variability could be a feature rather than a bug. Indeed, individual experiences could be replayed with very different temporal organization, and the identifiability of SWRs as content-full representations was similar across a wide range of clocks. At the same time, the events tended to be less classifiable, but even these event could serve a purpose. Identifiable events, by definition are those that contain sequential activation of adjacent spatial elements. These events, by themselves, might not be sufficient to drive associations among places that are in the same environment but not part sequential trajectories. We therefore suggest that the more random, non-identifiable events could serve to create broader associations across non-adjacent locations that could be useful to grouping experiences that occur within the same overall context. This would further suggest that SWR-associated replay is not a simple, uniform compression of experience, but rather an instantaneous, random sample (***Foster***, 2017; ***Stella et al***., 2019), and such a random sampling scheme allows for an unbiased, efficient representation of experience. This sampling hypothesis is also consistent with a recent study that showed hippocampal replay can represent Brownian diffusion-like random trajectories that cover position over wide ranges of spatiotemporal scales (***Stella et al***., 2019).

More broadly, human cognition is characterized by its extreme flexibility—the ability to transfer past learning to new contexts and to form abstract thoughts, such as analogies and inferences, to guide behaviors (***Behrens et al***., 2018). One crucial prerequisite of this flexibility is the ability to remember past experiences at different levels of specificity. In particular, advances in reinforcement learning showed that there are advantages to learning and memory when past experience is represented at different levels of temporal abstraction (***Schmidhuber***, 1992; ***Sutton et al***., 1999; ***Bakker and Schmidhuber***, 2004; ***Kulkarni et al***., 2016; ***Levy et al***., 2019; ***Konidaris***, 2019). More recently, a human imaging study observed that speed of time-compressed forward replay flexibly changes in human episodic memory (***Michelmann et al***., 2019). Our observations suggest that the hippocampal circuits replay spatial experiences at multiple levels on an event-to-event basis during SWRs. We propose that such a variable “clock” might constitute a general mechanism for flexible memory storage that could enable subsequent flexibility in behavioral choices.

## Acknowledgements

We thank Liam Paninski, Uri Eden, and members of the Eden group for helpful discussion. We thank Kenneth Kay for sharing his data on animal 3 and for insightful comments on earlier versions of the manuscript. This work began when X.D. was a postdoctoral research scientist in Liam Paninski’s group. X.D. additionally thanks Liam Paninski and the Center for Theoretical Neuroscience at Columbia University for organizational support during her time there. This work was supported by the National Natural Science Foundation of China (grant No.12001024 to X.D.), the NSF NeuroNex Award (DBI-1707398), the NIH (R01 MH-105174 to L.M.F.), the Gatsby Charitable Foundation, and the Simons Foundation (SCGB 521921 to L.M.F. and SCGB 543023).

## Author contributions

X.D. and L.M.F. conceived the study. M.S., and M.P.K. conducted the experiments. X.D. analyzed the data, with input from S.C., X-X.W., and L.M.F. All authors provided feedback. X.D. and L.M.F. wrote the paper.

## METHODS

### Subjects, neural recording, and behavioral task

The experimental methods are described in detail in ***Karlsson and Frank***, 2008, 2009; ***Kay et al***., 2016. In brief, data were taken from four male Long-Evans rats (500-600 g) implanted with a microdrive array containing 21, 30, 14, 30 independently movable tetrodes, respectively, targeting CA1 and CA3. Tetrodes that never yielded clusterable units across the 11, 10, 8, 12 days of experiments, respectively, were excluded from analysis. Following histological verification, tetrodes that ended up in areas other than CA1 and CA3 were excluded from analysis. In the analyses presented here, multiunit activity from 16, 18, 14, 17 simultaneously recorded tetrodes, respectively, were included. Unsorted spikes with peak-to-trough width of less than 0.35 ms were identified as from putative inhibitory interneurons, and unsorted spikes with peak-to-trough width of more than 0.35 ms were identified as from pyramidal neurons.

The multiunit activity (MUA) recorded from one tetrode consists of all detectable spike waveforms that cross a minimum amplitude threshold, usually set between 40 and 100 mV. Through standard manual clustering techniques, between 0 and 10 well-isolated units were extracted per tetrode in the data set described here. An average of 23, 53, 30, 45 single units across all recording epochs, respectively, were extracted. Importantly, the activity of these sorted putative units includes a minority of detectable spikes: An average of 92.94%, 89.13%, 94.80%, 91.12% of spikes across all recording epochs, respectively, remain unclassified because they could not be confidently assigned to a single unit. Thus, a large fraction of the data collected in these recordings would not be used for standard decoding analyses.

The hippocampal data in this paper are collected when the four animals were trained to perform a continuous alternation task on an W-maze (76 cm X 76 cm with 7-cm-wide track) shown in Figure 1A for liquid reward (condensed milk). Each task epoch lasted on average 15 minutes. The animal was rewarded each time it visited the end of an arm in the correct sequence, starting in the center and then alternating visits to each outer arm and returning to the center. The animal would start at the food well in position O and run toward the intersection, or “choice point” (position CP), at the top of the center stem, where a choice would need to be made. The correct choice is to alternate between left and right on successive turns. If a correct choice is made, for example, to turn right, the animal would continue to run toward the right food well (position W(right)) to receive a reward, and then return to the center well at position O to move on to the next trial. If an error is made, the rat is not given a reward and must return to the center well to initiate the next trial. All animal procedures and surgery were reviewed and approved by the University of California San Francisco Institutional Animal Care and Use Committee and were in accordance with National Institutes of Health guidelines.

We linearized the actual two-dimensional coordinate position of the rat to a single coordinate. The one-dimensional coordinate indicates the total distance from the center well (position O) in centimeters, with negative numbers indicating trajectories that include a left turn and positive numbers indicating trajectories that include a right turn. When the rat was on the center arm of the maze, the region to which its position was mapped was determined by the direction from which the rat came during inbound trajectories and by the direction it would turn next when it reached the choice point (position CP) during outbound trajectories. Throughout this report, to facilitate ease of visualization, when plotting we label the linearized maze at position O (“origin”), CP (“choice point”), or W (“Well”, left or right) instead of the corresponding signed one-dimensional coordinate.

### SWR detection

Ripples were detected on 4, 11, 7, 5 tetrodes recorded from CA1, respectively, by using an aggregated measure of the root mean square (RMS) power in the 150-250 Hz band across the tetrodes (***Csicsvari et al***., 1999). The aggregated RMS power was then smoothed with a kernel (4-ms standard deviation), and SWR events were detected as lasting at least 15 ms above 2 standard deviations of the mean. The entire SWR time was then set to include times immediately before and after the power exceeded the mean. SWR events included in this analysis were speed-thresholded at 4 cm/sec. Recording epochs with less than 50 detected, speed-thresholded SWR events were excluded from analysis; under this criterion, 3, 0, 0, 2 epochs, respectively, were excluded.

### Data analysis

All analyses were carried out using custom software written in MATLAB (MathWorks).

### Rhythmic organization of population spiking underlying individual SWRs

The temporal organization of population spiking underlying individual SWR event was defined using multiunit spiking activity. The population spiking activity was calculated by summing up unsorted spikes recorded on tetrodes that had yielded clusterable units across all days of experiments. We then computed the autocorrelation function of this summed spike train binned by 1 ms with lags up to 200 ms. When sorting spiking autocorrelation across events (e.g. Figure 2), we first smoothed the autocorrelation function with a 20ms Gaussian kernel and then calculated the lags of the first positive side peak. We found the peak lags remained consistent for a range of smoothing window between 10 ms and 20 ms.

### Rhythmic organization of population spiking underlying individual RUN lap

The temporal organization of population spiking underlying individual RUN lap was defined using multiunit spiking activity. The population spiking activity was calculated by summing up unsorted spikes recorded on tetrodes that had yielded clusterable units across all days of experiments. We then computed the autocorrelation of this summed spike train binned by 1 ms with lags up to 400 ms. When sorting spiking autocorrelation across events (e.g. Figure 2), we first smoothed the autocorrelation function with a 100ms Gaussian kernel and then calculated the lags of the first positive side peak.

### Rhythmic organization of population spiking underlying individual theta cycles

We first computed an LFP-defined theta cycles: LFP was filtered at 5-11 Hz. Peaks and troughs of the filtered LFP were detected and used to define half-cycles by linear interpolation. Theta cycles were identified as individual cycles whose duration was consistent with the 5-11 Hz frequency range (<200 ms (5 Hz) and >90 ms (11 Hz)). We then computed an MUA-defined theta cycles: The population spiking activity was calculated by summing up unsorted spikes recorded on tetrodes that had yielded clusterable units across all days of experiments. This MUA was smoothed with a 100 msec Gaussian windows. Peaks and troughs of the smoothed MUA were detected. MUA-defind theta cycles were identified as time windows between troughs. Theta cycles included in the analyses for this manuscript were defined by a consensus measure between LFP and multiunit activity: if an LFP-defined theta cycle and an MUA-defined theta cycle overlap with less than +/-25 msec (both for start and end of a cycle) discrepancy, it was included for subsequent analyses. The temporal organization of population spiking underlying individual theta cycles was defined using multiunit spiking activity, similar to SWRs: The population spiking activity was calculated by summing up unsorted spikes recorded on tetrodes that had yielded clusterable units across all days of experiments. We then computed the autocorrelation function of this summed spike train binned by 1 ms with lags up to 200 ms. When sorting spiking autocorrelation across events (e.g. Figure 2), we first smoothed the autocorrelation function with a 20ms Gaussian moving window and then calculated the lags of the first positive side peak. Only theta cycles whose peak lags remained consistent for a range of smoothing window between 15 ms and 20 ms were included in the analysis.

### Parameter estimation for the distributional fit

We first computed the histogram of the timings of the first positive side peak lag of SWR-associated population spiking autocorrelations across events. We then fit lognormal distribution to this histogram whose parameters, mu and sigma, were estimated with maximum likelihood.

### Clusterless decoding of representational content

We decoded the spatial representation content of individual SWR event using a clusterless decoding method (***Deng et al***., 2015) that does not require multiunit spiking waveforms to be sorted into single units and instead incorporates waveform information of unsorted spikes, by using the theory of marked point process: Any point process representing neural spiking can be fully characterized by its conditional intensity function. A conditional intensity function describes the instantaneous probability of observing a spike, given previous spiking history. By relating the conditional intensity to specific biological and behavioral signals we can specify a spike train encoding model. The conditional intensity also generalizes to the marked case, in which a random vector, termed a mark, is attached to each point. Here we use the mark to characterize features of the spike waveform. In the case of tetrode recordings, the mark is a length-four vector of the maximum amplitudes on each of the four electrodes at every spike time.

Briefly, we characterizes the instantaneous probability of observing a spike with mark 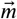 at time *t* as a function of some underlying internal state variable *x*(*t*), such as an animal’s location in space, that varies across time, using the joint mark intensity function 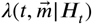 where *H_t_* is the history of the spiking activity up to time *t*:

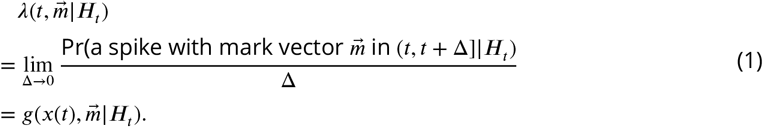

Our decoding algorithm, using discrete-time state-space adaptive filters, computes, at each time point, the unnormalized posterior distribution of the state variable given observed marked spiking activity:

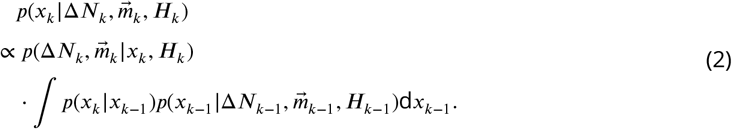

The 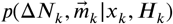 term is the likelihood or observation distribution at the current time:

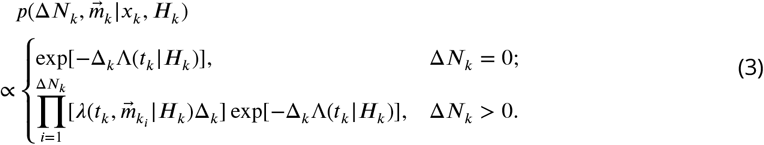

The 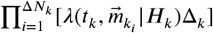 term characterizes the distribution of firing Δ*N_k_* spikes, such that the mark value of the *i*th spike in the interval (*t*_*k*−1_, *t_k_*] is *m_k_i__*, where *i* = 1,…, Δ*N_k_*. The probability of observing a spike regardless of the mark values is denoted by 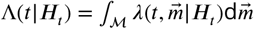.

### Classification of representational content

We extended the state variable in the clusterless decoder to jointly include a discrete decision state, *I*, which identifies whether each SWR event represents an outbound or inbound trajectory as well as whether the temporal order of activity occurs forward or backward in time (***Deng et al***., 2016). In our analyses, *I* is an indicator function for a replay event being one of the following four categories: “outbound, forward”, “outbound, reverse”, “inbound, forward”, or “inbound, reverse”. We categorized the representational content of example replay events using a marked point process filter with the joint state variable *x*(*t*), *I*. Finally, if the posterior probability of the decision state, Pr(*I*), of an individual SWR event, first passes a threshold of 98% for consecutive 10 temporal bins for a particular category of *I*, we assigned the event to that category. SWR events whose decision state probabilities did not pass the threshold and whose representational content cannot be classified into one of the four categories were assigned to a fifth category as “unclassified”.

### SWR-triggered spectrogram and local field frequency measures

SWR-triggered spectrograms were computed using the multitaper method and a 100 ms sliding windows with a 10 ms step size. A z-score was computed for each frequency band using the mean and standard deviation of the power calculated across the entire behavioral session for each tetrode. For each 100 ms bin, we obtained a normalized measure of power for each frequency band (sharp wave: < 20 Hz; slow gamma: 20-50 Hz; ripple: 150-250 Hz) in units of standard deviation from the mean. To quantify gamma phase locking during SWRs, the phase of coherence for the gamma band was averaged across all CA3-CA1 tetrode pairs for each SWR. Thus, each SWR contributed a single value for each 100 ms temporal bin relative to SWR detection. We combined values across SWRs to obtain a distribution of gamma phase offsets in each bin. The angular variance of this distribution was taken as a measure of phase locking for each epoch.

### Hypothesis testing

Bootstrap methods (1000 iterations) were used to test homogeneity of variances between peak lags of two behavioral states, Wilcoxon rank-sum tests were used when comparing between two groups of peak lags of spiking autocorrelation within the same epoch, Kruskal-Wallis tests were used when comparing across multiple groups, and Student’s t-tests were used when comparing the mean of a variable across all lags with zero. Across all epochs for an individual animal, summary of p-values was reported either in text or in supplementary figures. Rayleigh’s test for detecting unimodal deviation from circular uniformity was used to evaluate the phase-locking of CA3 multiunit spiking to CA1 slow gamma. To evaluate the effects of multiple covariates on the peak lag of the spiking autocorrelation across all epochs for an individual animal, linear mixed-effects models were used to account for random effects of epoch. Student’s t-tests were used to estimate the statistical significance of individual fixed effects coefficients. For better interpretable units, standardized coefficients for discrete independent covariates were computed by centering; standardized coefficients for continuous independent covariates are computed by first centering and then dividing by two standard deviations (***Gelman***, 2008).

---

**Figure S1.**
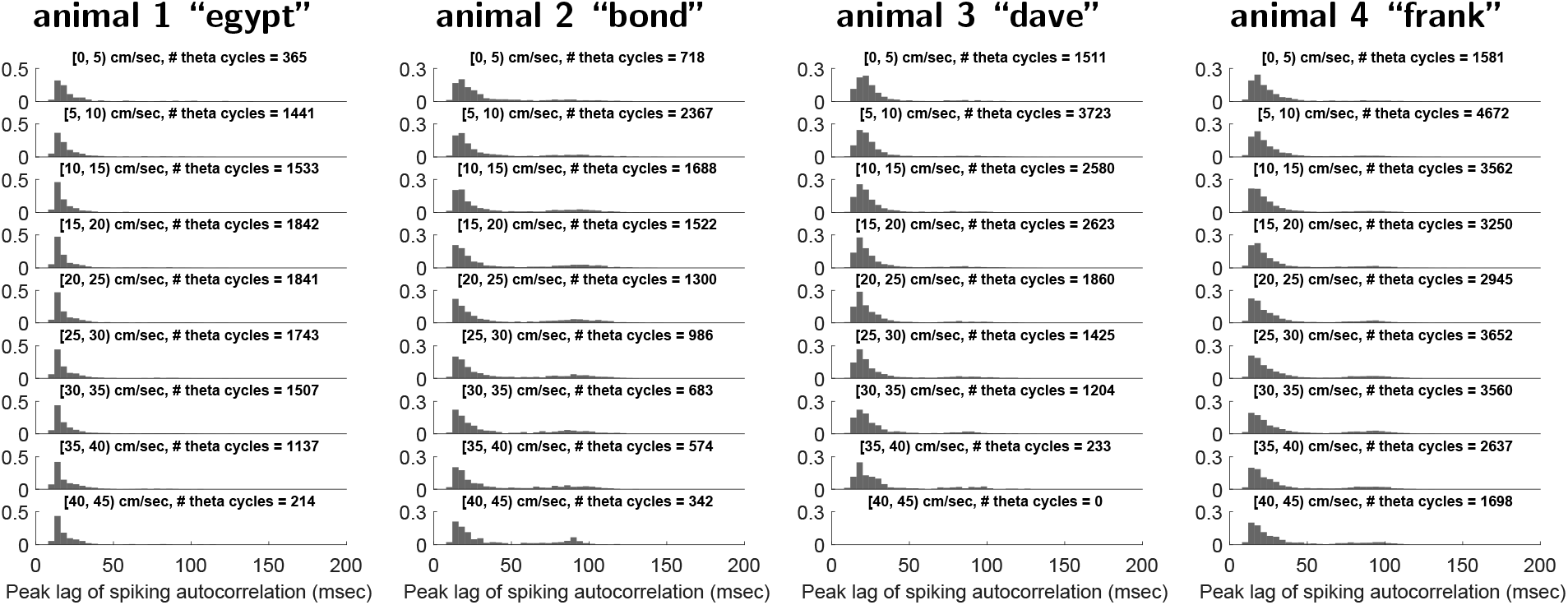
Theta cycles with lower speeds were more likely to be associated with larger mode values of the empirical distributions. Each panel shows the relative frequency histogram of the first side peak of the theta cycle spiking autocorrelation within a speed group for one animal. Correlation coefficient between the mode of the empirical distribution and speed: animal 1, −0.70 (0.04); animal 2, −0.78 (0.01); animal 3, −0.69 (0.06); animal 4, −0.46 (0.22).

**Figure S2.**
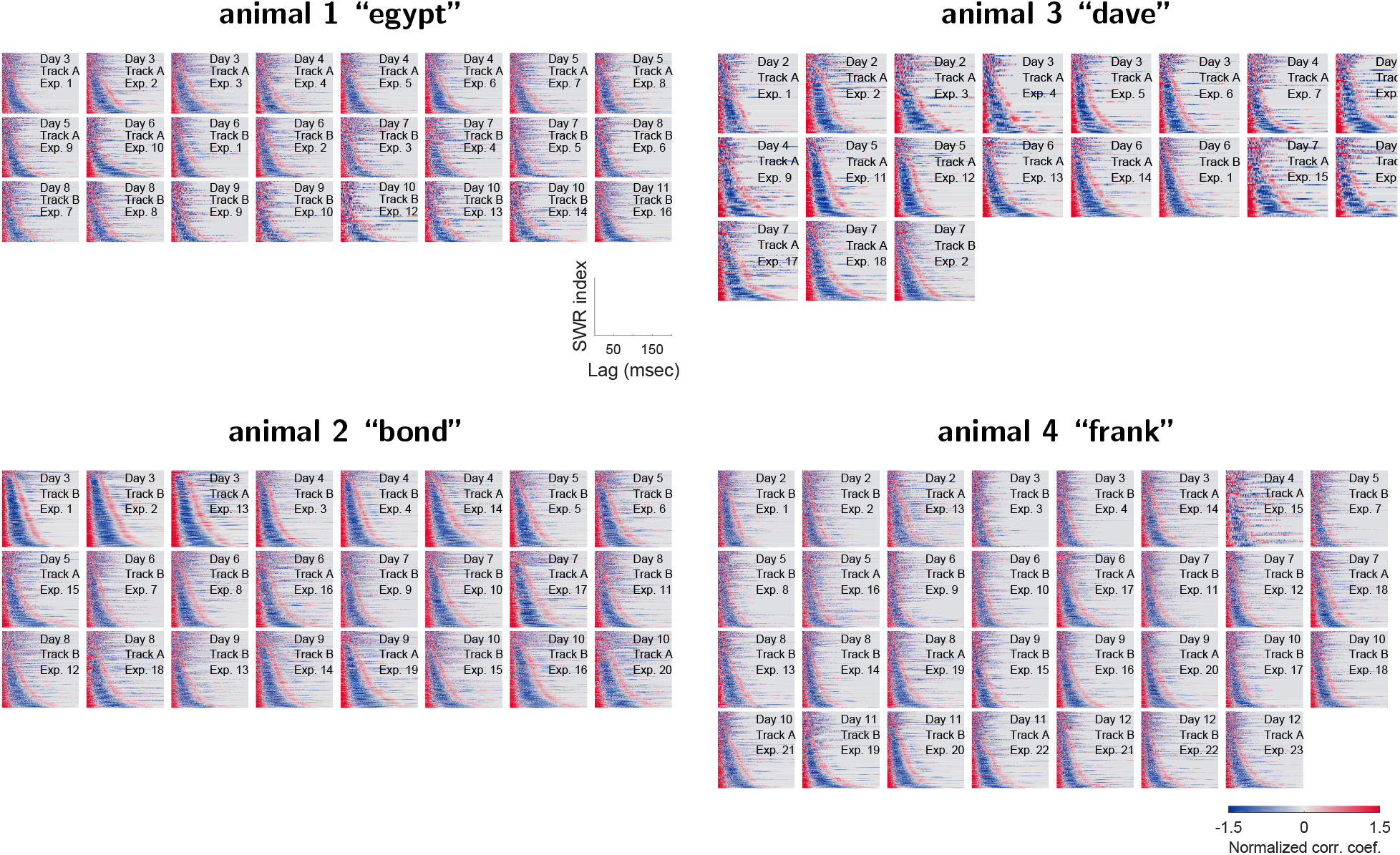
Event-to-event variability in SWR-associated spiking organization across recording epochs, across days, and across animals (related to Figure 1). Each panel plots the normalized autocorrelations of SWR-associated multiunit spiking activity in an individual epoch, collated in ascending order by the timings of the first positive side peak lag. Every horizontal line is the autocorrelation function of an individual SWR, respectively; only non-negative lags are displayed. Positive and negative correlation coefficients are plotted in shades of red and blue, respectively. The overlay text on each panel denotes the experimental day, the track ID and the number of exposure for that particular epoch.

**Figure S3.**
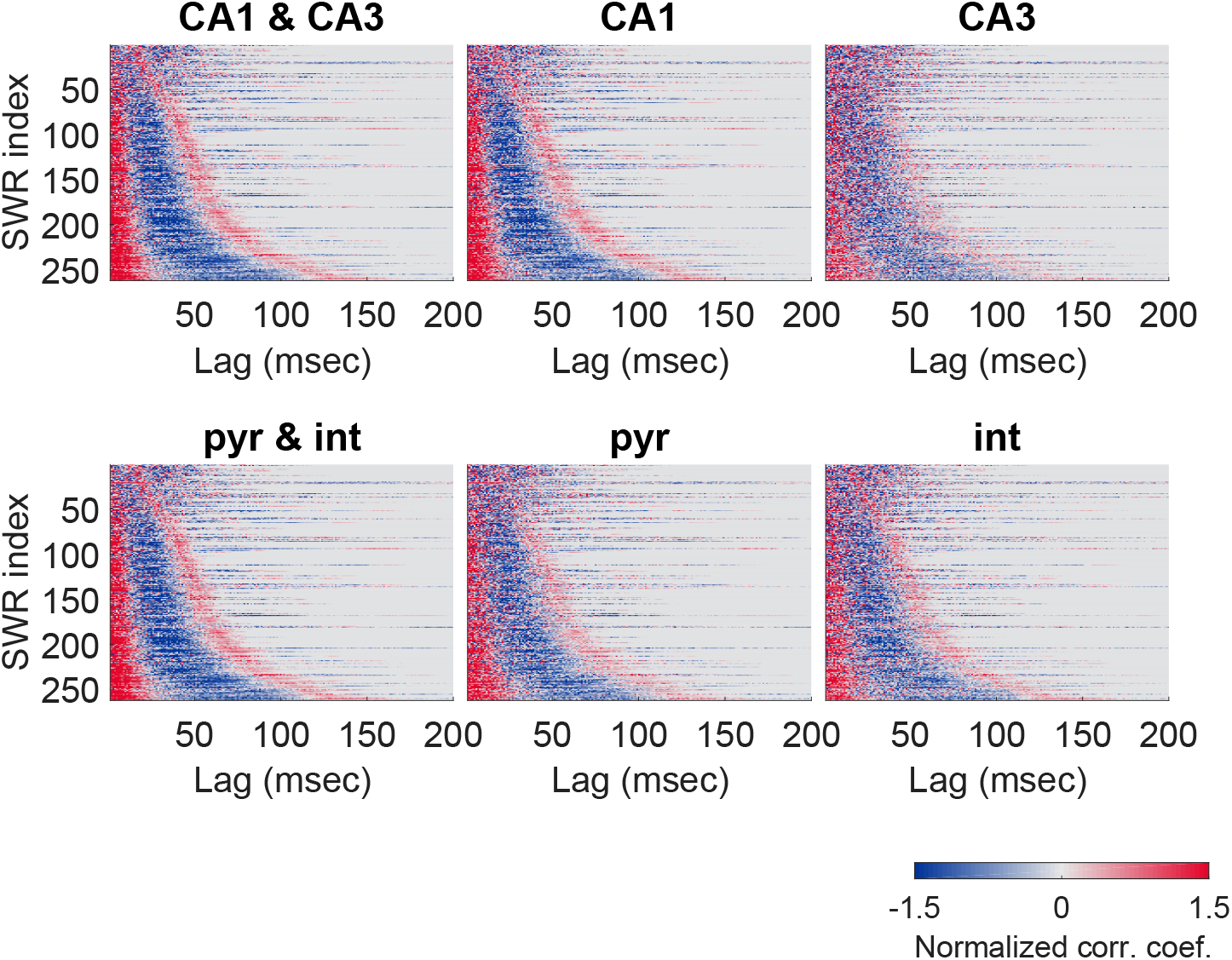
SWR-associated rhythmic population activity in CA3 is noisier than in CA1, but appeared to be similar between pyramidal cell populations and interneuron populations (related to Figure 1).) Each panel plots the normalized autocorrelations of SWR-associated multiunit spiking activity in an example epoch across hippocampal subfields (*top panels*) or across cell types (*bottom panels*), collated in ascending order by the timings of the first positive side peak lag. Every horizontal line is the autocorrelation function of an individual SWR; only non-negative lags are displayed. Positive and negative correlation coefficients are plotted in shades of red and blue, respectively.

**Figure S4.**
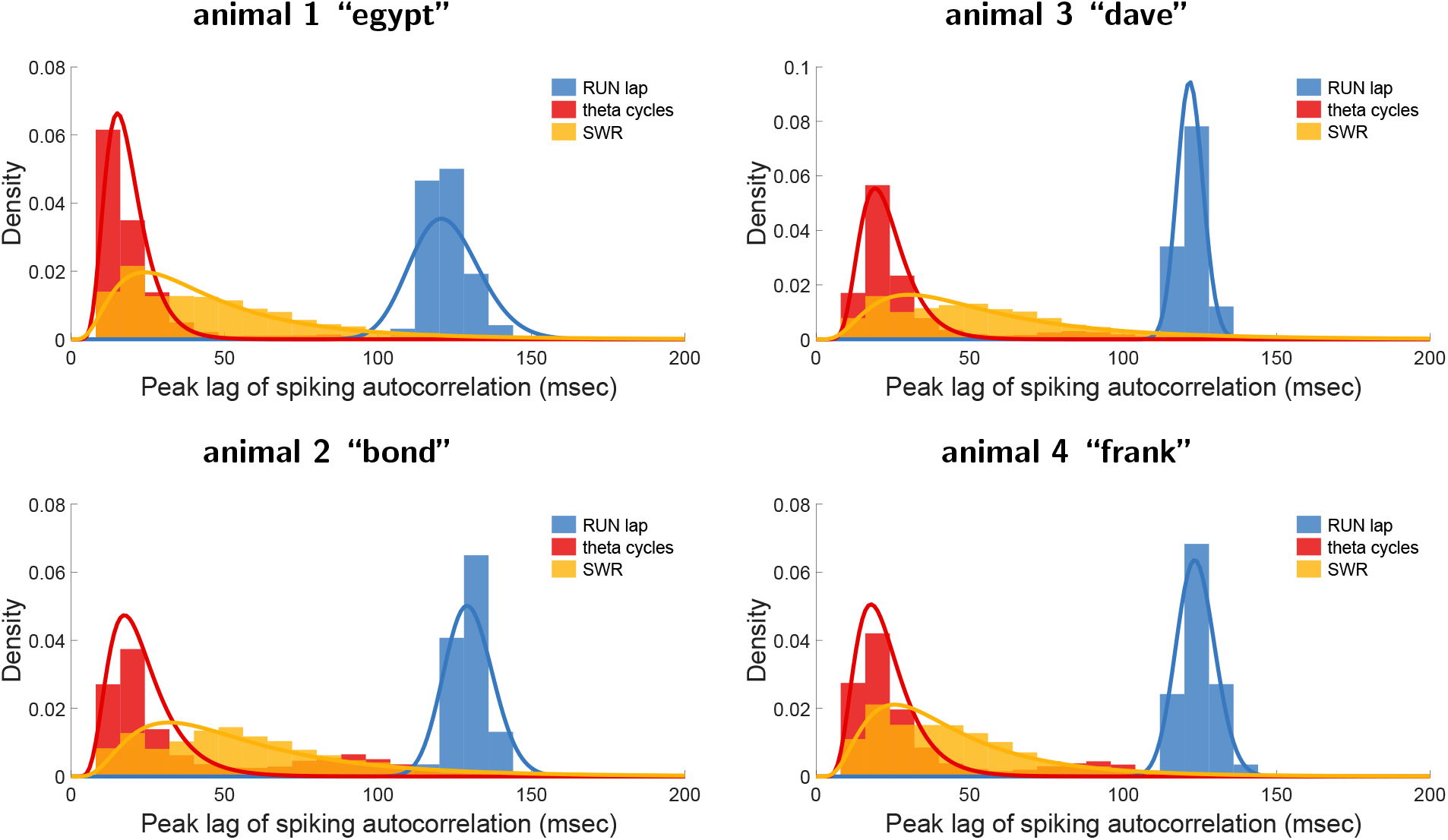
Periodicity of SWR-associated spiking organization follows an approximate lognormal distribution across animals (related to Figure 2). Each panel plots the histogram of the timing of the first positive side peak lag of RUN lap (blue), theta-cycle (red), and SWR (yellow) autocorrelation function, for an example animal. Lognormal distribution fitted to the gamma range of each histogram is plotted as solid line.

**Figure S5.**
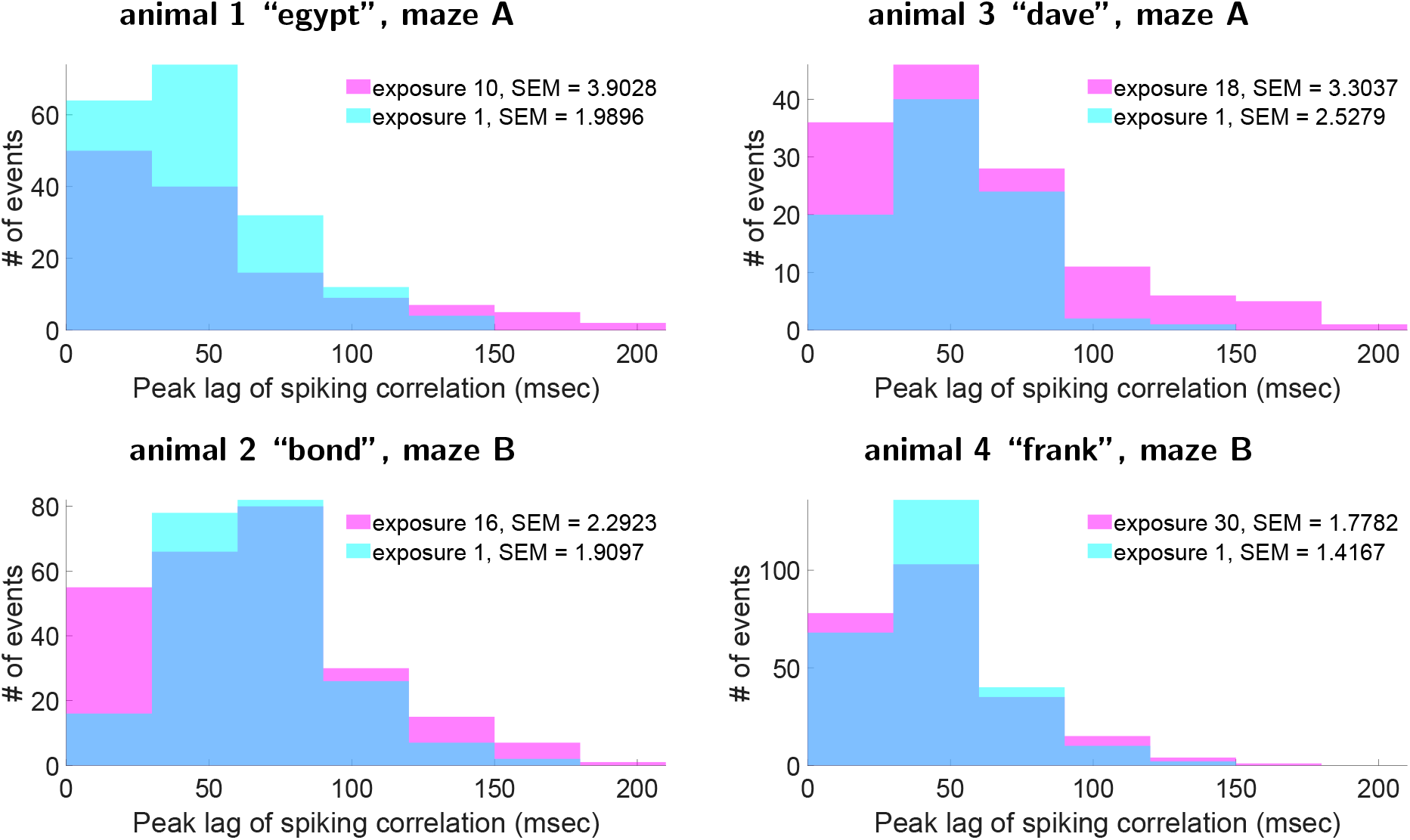
Empirical distribution of SWR slow gamma clock speed increased in speadth between the first and last task exposure (related to Figure 2). Each panel plots the histogram of the timing of SWR autocorrelation function’s first positive side peak lag for an example maze.

**Figure S6.**
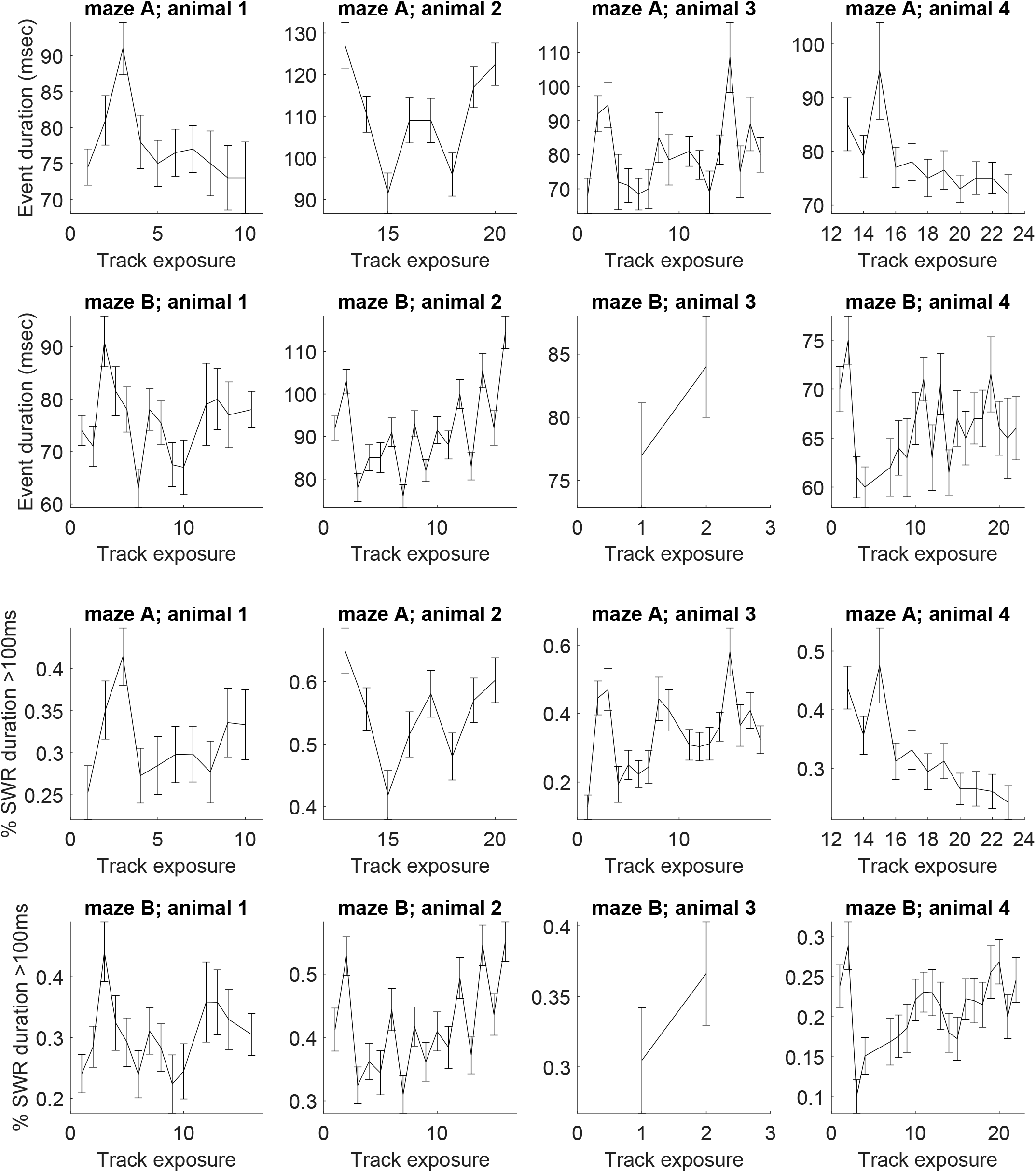
SWR event duration across mazes and across animals (related to Figure 2). (First and second rows), SWR event duration across task exposures (median ±standard error). (Third and Last rows), proportion of long SWRs across task exposures (mean ±standard error).

## References

Atallah BV, Scanziani M. Instantaneous Modulation of Gamma Oscillation Frequency by Balancing Excitation with Inhibition. Neuron. 2009; 62:566–577. doi: 10.1016/j.neuron.2009.04.027.

Bakker B, Schmidhuber J. Hierarchical reinforcement learning based on subgoal discovery and subpolicy specialization. In: Proceedings of the 8th Conference on Intelligent Autonomous Systems; 2004. p. 438–445.

Behrens TEJ, Muller TH, Whittington JCR, Mark S, Baram BA, Stachenfeld KL, Kurth-Nelson Z. What Is a Cognitive Map? Organizing Knowledge for Flexible Behavior. Neuron. 2018; 100:490–509. doi: 10.1016/j.neuron.2018.10.002.

Bragin A, Jando G, Nadasday Z, Hetke J, Wise K, Buzsaki G. Gamma (40-100 Hz) oscillation in the hippocampus of the behaving rat. Journal of Neuroscience. 1995; 15(1):47–60. doi: 10.1523/jNEUROSCI.18-01-00388.1998.

Buzsaki G, Logothetis N, Singer W. Scaling brain size, keeping timing: Evolutionary preservation of brain rhythms. Neuron. 2013; 80:751–764. doi: 10.1016/j.neuron.2013.10.002.

Buzsaki G. Two-stage model of memory trace formation: A role for “noisy” brain states. Neurosci. 1989; 31:551–570. doi: 10.1016/0306-4522(89)90423-5.

Buzsaki G. Theta oscillations in the hippocampus. Neuron. 2002; 33:325–340. doi: 10.1016/s0896-6273(02)00586-x.

Buzsaki G. Hippocampal sharp wave-ripple: A cognitive biomarker for episodic memory and planning. Hippocampus. 2015; 25:1083–1188. doi: 10.1002/hipo.22488.

Buzsaki G. The Brain From Inside Out. New York, NY: Oxford University Press; 2019. doi: 10.1093/oso/9780190905385.001.0001.

Carr MF, Karlsson MP, Frank LM. Transient slow gamma synchrony underlies hippocampal memory replay. Neuron. 2012; 75:700–713. doi: 10.1016/j.neuron.2012.06.014.

Cheng S, Frank LM. New Experiences Enhance Coordinated Neural Activity in the Hippocampus. Neuron. 2008; 57:303–313. doi: 10.1016/j.neuron.2007.11.035.

Colgin LL, Denninger T, Fyhn M, Hafting T, Bonnevie T, Jensen O, Moser MB, Moser EI. Frequency of gamma oscillations routes flow of information in the hippocampus. Nature. 2009; 462:353–357. doi: 10.1038/na-ture08573.

Csicsvari J, Hirase H, Czurko A, Mamiya A, Buzsaki G. Oscillatory coupling of hippocampal pyramidal cells and interneurons in the behaving rat. J Neurosci. 1999; 19:274–287. doi: 10.1523/JNEUROSCI.19-01-00274.1999.

Csicsvari J, Hirase H, Mamiya A, Buzsaki G. Ensemble patterns of hippocampal CA3-CA1 neurons during sharp wave-associated population events. Neuron. 2000; 28:585–594. doi: 10.1016/s0896-6273(00)00135-5.

Davidson TJ, Kloosterman F, Wilson MA. Hippocampal replay of extended experience. Neuron. 2009; 63:497–507. doi: 10.1016/j.neuron.2009.07.027.

Deng X, Liu DF, Karlsson MP, Frank LM, Eden UT. Rapid classification of hippocampal replay content for real-time applications. J Neurophysiol. 2016; 116:2221–2235. doi: 10.1152/jn.00151.2016.

Deng X, Liu DF, Kay K, Frank LM, Eden UT. Clusterless decoding of position from multiunit activity using a marked point process filter. Neural Comput. 2015; 27:1438–1460. doi: 10.1162/NECO_a_00744.

Eichenbaum H, Cohen NJ. From Conditioning to Conscious Recollection: Memory systems of the brain. New York, NY: Oxford University Press; 2004. doi: 10.1093/acprof:oso/9780195178043.001.0001.

Fernandez-Ruiz A, Oliva A, de Oliveira EF, Rocha-Almeida F, Tingley D, Buzsaki G. Long-duration hippocampal sharp wave ripples improve memory. Science. 2019; 364:1082–1086. doi: 10.1126/science.aax0758.

Foster DJ. Replay comes of age. Annu Rev Neurosci. 2017; 40:581–602. doi: 10.1146/annurev-neuro-072116-031538.

Foster DJ, Wilson MA. Reverse replay of behavioural sequences in hippocampal place cells during the awake state. Nature. 2006; 440:680–683. doi: 10.1038/nature04587.

Gelman A. Scaling regression inputs by dividing by two standard deviations. Statist Med. 2008; 27:2865–2873. doi: 10.1002/SIM.3107.

Gillespie AK, Astudillo Maya DA, Denovellis EL, Liu DF, Kastner DB, Coulter ME, Roumis DK, Eden UT, Frank LM. Hippocampal replay reflects specific past experiences rather than a plan for subsequent choice. bioRxiv. 2021; doi: 10.1101/2021.03.09.434621.

Gillespie AK, Jones EA, Lin YH, Karlsson MP, Kay K, Yoon SY, Tong LM, Nova P, Carr JS, Frank LM, Huang Y. Apolipoprotein E4 causes age-dependent disruption of slow gamma oscillations during hippocampal sharp-wave ripples. Neuron. 2016; 90:740–751. doi: 10.1016/j.neuron.2016.04.009.

Gupta AS, van der Meer MAA, Touretzky DS, Redish AD. Segmentation of spatial experience by hippocampal theta sequences. Nat Neurosci. 2012; 15:1032–1039. doi: 10.1038/nn.3138.

Hemberger M, Shein-Idelson M, Pammer L, Laurent G. Reliable Sequential Activation of Neural Assemblies by Single Pyramidal Cells in a Three-Layered Cortex. Neuron. 2019; 104:353–369. doi: 10.1016/j.neuron.2019.07.017.

Hinman JR, Penley SC, Long LL, Escabi MA, Chrobak JJ. Septotemporal variation in dynamics of theta: speed and habituation. J Neurophysiol. 2011; 105:2675–2686. doi: 10.1152/jn.00837.2010.

Joo HR, Frank LM. The hippocampal sharp wave-ripple in memory retrieval for immediate use and consolidation. Nat Rev Neurosci. 2018; 19:744–757. doi: 10.1038/s41583-018-0077-1.

Karlsson MP, Frank LM. Network dynamics underlying the formation of sparse, informative representations in the hippocampus. J Neurosci. 2008; 28:14271–14281. doi: 10.1523/JNEUROSCI.4261-08.2008.

Karlsson MP, Frank LM. Awake replay of remote experiences in the hippocampus. Nat Neurosci. 2009; 12:913–918. doi: 10.1038/nn.2344.

Kay K, Frank LM. Three brain states in the hippocampus and cortex. Hippocampus. 2019; 29:184–238. doi: 10.1002/hipo.22956.

Kay K, Sosa M, Chung JE, Karlsson MP, Larkin MC, Frank LM. A hippocampal network for spatial coding during immobility and sleep. Nature. 2016; 531(7593):185–190. doi: 10.1038/nature17144.

Kemere C, Carr MF, Karlsson M, Frank LM. Rapid and Continuous Modulation of Hippocampal Network State during Exploration of New Places. PLos One. 2013; 8:e73114. doi: 10.1371/journal.pone.0073114.

Kloosterman F, Layton SP, Chen Z, Wilson MA. Bayesian decoding using unsorted spikes in the rat hippocampus. J Neurophysiol. 2014; 111:217–227. doi: 10.1152/jn.01046.2012.

Konidaris G. On the necessity of abstraction. Curr Opin Behav Sci. 2019; 29:1–7. doi: 10.1016/j.cobeha.2018.11.005.

Kulkarni TD, Narasimhan K, Saeedi A, Tenenbaum J. Hierarchical Deep Reinforcement Learning: Integrating Temporal Abstraction and Intrinsic Motivation. In: Advances in Neural Information Processing Systems; 2016. p. 3682–3690.

Lee AK, Wilson MA. Memory of sequential experience in the hippocampus during slow wave sleep. Neuron. 2002; 36:1183–1194. doi: 10.1016/s0896-6273(02)01096-6.

Levy A, Konidaris G, Platt R, Saenko K. Learning Multi-Level Hierarchies with Hindsight. In: Proceedings of the 7th International Conference on Learning Representations; 2019..

Lisman J. The theta/gamma discrete phase code occuring during the hippocampal phase precession may be a more general brain coding scheme. Hippocampus. 2005; 15:913–922. doi: 10.1002/hipo.20121.

Lisman J, Buzsaki G. A neural coding scheme formed by the combined function of gamma and theta oscillations. Schizophrenia Bulletin. 2008; 34(5):974–980. doi: 10.1093/schbul/sbn060.

Lisman JE, Jensen O. The theta-gamma neural code. Neuron. 2013; 77:1002–1016. doi: 10.1016/j.neuron.2013.03.007.

Lopes-Dos-Santos V, van de Ven GM, Morley A, Trouche S, Campo-Urriza N, Dupret D. Parsing hippocampal theta oscillations by nested spectral components during spatial exploration and memory-guided behavior. Neuron. 2018; 100:940–952. doi: 10.1016/j.neuron.2018.09.031.

Michelmann S, Staresina BP, Bowman H, Hanslmayr S. Speed of time-compressed forward replay flexibly changes in human episodic memory. Nat Hum Behav. 2019; 3:143–154. doi: 10.1038/s41562-018-0491-4.

O’Keefe J, Dostrovsky J. The hippocampus as a spatial map. Preliminary evidence from unit activity in the freely-moving rat. Brain Res. 1971; 34:171–175. doi: 10.1016/0006-8993(71)90358-1.

Oliva A, Fernandez-Ruiz A, de Oliveira EF, Buzsaki G. Origin of Gamma Frequency Power during Hippocampal Sharp-Wave Ripples. Cell Reports. 2018; 25:1693–1700. doi: 10.1016/j.celrep.2018.10.066.

O’Neill J, Senior T, Csicsvari J. Place-selective firing of CA1 pyramidal cells during sharp wave/ripple network patterns in exploratory behavior. Neuron. 2006; 49:143–155. doi: 10.1016/j.neuron.2005.10.037.

Pfeiffer BE, Foster DJ. Autoassociative dynamics in the generation of sequences of hippocampal place cells. Science. 2015; 349:180–183. doi: 10.1126/science.aaa9633.

Schmidhuber J. Learning complex, extended sequences using the principle of history compression. Neural Computation. 1992; 4:234–242. doi: 10.1162/neco.1992.4.2.234.

Sirota A, Csicsvari J, Buhl D, Buzsaki G. Communication between neocortex and hippocampus during sleep in rodents. Proc Natl Acad Sci USA. 2003; 100:2065–2069. doi: 10.1073/pnas.0437938100.

Skaggs WE, McNaughton BL, Wilson MA, Barnes CA. Theta phase precession in hippocampal neuronal populations and the compression of temporal sequences. Hippocampus. 1996; 6:149–172. doi: 10.1002/(SICI)1098-1063(1996)6:2<149::AID-HIPO6>3.0.CO;2-K.

Stella F, Baracskay P, O’Neill J, Csicsvari J. Hippocampal Reactivation of Random Trajectories Resembling Brownian Diffusion. Neuron. 2019; 102:1–12. doi: 10.1016/j.neuron.2019.01.052.

Sutton RS, Precup D, Singh S. Between MDPs and semi-MDPs: A framework for temporal abstraction in reinforcement learning. Artificial Intelligence. 1999; 112:181–211. doi: 10.1016/S0004-3702(99)00052-1.

